# Haplotype sharing provides insights into fine-scale population history and disease in Finland

**DOI:** 10.1101/200113

**Authors:** Alicia R. Martin, Konrad J. Karczewski, Sini Kerminen, Mitja Kurki, Antti-Pekka Sarin, Mykyta Artomov, Johan G. Eriksson, Tõnu Esko, Giulio Genovese, Aki S. Havulinna, Jaakko Kaprio, Alexandra Konradi, László Korányi, Anna Kostareva, Minna Männikkö, Andres Metspalu, Markus Perola, Rashmi B. Prasad, Olli Raitakari, Oxana Rotar, Veikko Salomaa, Leif Groop, Aarno Palotie, Benjamin M. Neale, Samuli Ripatti, Matti Pirinen, Mark J. Daly

## Abstract

Finland provides unique opportunities to investigate population and medical genomics because of its adoption of unified national electronic health records, detailed historical and birth records, and serial population bottlenecks. We assemble a comprehensive view of recent population history (≤100 generations), the timespan during which most rare disease-causing alleles arose, by comparing pairwise haplotype sharing from 43,254 Finns to geographically and linguistically adjacent countries with different population histories, including 16,060 Swedes, Estonians, Russians, and Hungarians. We find much more extensive sharing in Finns, with at least one ≥ 5 cM tract on average between pairs of unrelated individuals. By coupling haplotype sharing with fine-scale birth records from over 25,000 individuals, we find that while haplotype sharing broadly decays with geographical distance, there are pockets of excess haplotype sharing; individuals from northeast Finland share several-fold more of their genome in identity-by-descent (IBD) segments than individuals from southwest regions containing the major cities of Helsinki and Turku. We estimate recent effective population size changes over time across regions of Finland and find significant differences between the Early and Late Settlement Regions as expected; however, our results indicate more continuous gene flow than previously indicated as Finns migrated towards the northernmost Lapland region. Lastly, we show that haplotype sharing is locally enriched among pairs of individuals sharing rare alleles by an order of magnitude, especially among pairs sharing rare disease causing variants. Our work provides a general framework for using haplotype sharing to reconstruct an integrative view of recent population history and gain insight into the evolutionary origins of rare variants contributing to disease.

## Background

A central goal in human genetics is to identify causal disease variants, elucidate their functional roles, and pinpoint therapeutic routes for correction. Recent large-scale DNA sequencing consortia efforts such as ExAC have demonstrated that one of the most predictive features of pathogenicity is allele frequency, with most disease-causing variants being rare and thus relatively young ^1,2^. These variants have not yet been fully exposed to the forces of natural selection that common, older variants have survived. Genome-wide association studies (GWAS) continue to identify a myriad of common and increasingly rare risk variants across many traits and increase heritable variance explained ^3^, but their power is substantially reduced for rare variants. Additionally, standard GWAS approaches, such as the inclusion of principal components in GWAS to correct for population structure, are insufficient for rare variants ^4^.

Most rare variants that play a critical role in disease today arose during approximately the last 100 generations ^5^. Aside from *de novo* variants in early-onset developmental phenotypes, the role of recently evolved, large-effect variants in common disease is largely uncharacterized. Stronger effects are likely not confined to *de novo* variants but may persist for several generations; however, this class of variation has been difficult to distinguish with single variant analyses because of extremely limited power, especially for scenarios involving incomplete penetrance ^6,7^. It is imperative that we better understand recent population genetic history in this context because it bounds the ability of negative selection to purge deleterious variants, and is the most relevant period for producing disease-conferring variants subject to negative selection ^8,9^. Haplotype-based methods have two major benefits over single variant approaches for inferences into demographic history and disease association: 1) as opposed to commonly used site-frequency based approaches ^10^, they are more informative of population history during the last tens to hundreds of generations ago, and 2) they can expose disease-causing rare variants at the population level without necessitating deep whole-genome sequencing. Rather, haplotype sharing can take advantage of massive, readily available GWAS array data. While these advantages have been theoretically recognized when sample sizes were relatively small ^11,12^, they have been underutilized in the modern genomics era.

Finland provides a convenient example in which to infer both population history and rare disease associations because of unified electronic health records as well as the founder effect elicited by serial population bottlenecks. In addition to the out-of-Africa bottleneck experienced by all of Europe, Finland underwent multiple additional bottlenecks over the last few thousand years, with the Finnish founder population size estimated to include 3,000-24,000 individuals ^13–17^. Archaeological evidence indicates that Finland has been continuously inhabited since the end of the last ice age ~10.9kya, with a small population of not more than a few thousand early hunter-gatherers first settling throughout Finland mostly from the south and to a lesser extent from the east and a western Norwegian coastal route ^18^. A cultural split circa 2300 BC was hypothesized to separate the western and eastern areas of Finland, termed the Early and Late Settlement Regions (ESR and LSR), upon the arrival of the Corded Ware culture primarily restricted to the southwestern and coastal regions of the country; this split has been supported by Y chromosome and mitochondrial DNA as well as historical data ^18,19,20^. Archaeologists agree that Finland has historically been sparsely inhabited, but that the ESR encompassing the southern and western colonized regions of Finland was more densely and permanently settled beginning ~4,000 years ago. In contrast, the LSR encompassing the northern and eastern regions of Finland, were more permanently inhabited beginning in the 1500s, pushing existing nomadic Sami people further north into Lapland. Archaeological records suggest that a series of founding, extinction, and re-colonization events took place over two millennia before continuous habitation coincident with agriculture ^21^. While Finland was a part of the Swedish Kingdom until 1809 and then became a semi-autonomous grand duchy controlled by tsarist Russia until it gained independence in 1917, immigration into western and especially eastern Finland has been relatively low until after the collapse of the Soviet Union. Linguistically, roughly 5% of the population speaks Swedish as their mother tongue, and both Finnish and Swedish are taught at school. Bilingual Finns who speak Swedish as their mother tongue live mostly in the ESR in restricted western and southern coastal regions.

Because of serial bottlenecks in Finland, the site-frequency spectrum is skewed towards more common variants than other European populations, and deleterious alleles are more likely to be found in a homozygous state ^17^. The consequence of this is exemplified in the Finnish Disease Heritage (FinDis) database, which contains 36 monogenic diseases to date that are much more frequent in Finns than in any other population ^22^. Several complex diseases also show strong regional clines within Finland, for example with schizophrenia and familial hypercholesterolemia risk being greatest in northeastern Finland ^23,24^. Current Finnish demographic models are primarily based on single locus markers (i.e. the Y-chromosome and mitochondria) ^13,19,20^, and a few studies have recently expanded to incorporate autosomal data ^25–27^. In contrast to site frequency spectrum-based methods which consider sites independently and are therefore optimally powered to infer old demographic events (>100 generations ago), haplotype-based demographic inference is best powered to inform population history during the period most relevant for negatively selected traits (last 100 generations) ^28–31^. Multiple lines of evidence indicate that recent history is particularly important for disadvantageous traits. For example, long runs of homozygosity (ROH), a special case of recent haplotype sharing, are enriched for deleterious variation ^32^, and increased ROH have been associated with decreased educational attainment as well as intellectual disability ^33,34^. Further, allele dating techniques indicate that pathogenic variants are on average considerably younger than neutral variants ^2^.

In this study, we combine biobank-scale genetic and detailed birth record data to assemble a comprehensive inquiry into recent population history by employing genetic data from 43,254 Finnish individuals (~0.8% of Finland’s total population) and 16,060 demographically distinct individuals from geographically or linguistically neighboring countries, including Swedes, Estonians, Russians, and Hungarians. While Finland is a poised example for population insights from haplotype sharing due to serial population bottlenecks, our approach provides a general framework for using haplotype sharing to reconstruct an integrative view of recent population history (e.g. elucidation of migration, divergence, and population size changes over time) within and across countries. Through these analyses, we also demonstrate that elevated haplotype sharing patterns resulting from multiple population bottlenecks provide insights into the origins of certain genetic diseases.

## Results

### Population substructure across regions of Finland

To investigate fine-scale population structure within Finland, we assembled a panel of 43,254 Finnish individuals (**Table S1**, Methods). We performed principal components analysis (PCA) on all individuals, and using the subset of individuals with recorded birth record data show that genetic variation in Finland broadly reflects geographical birthplace (Methods), with highly significant correlations between PC1 and longitude (ρ=−0.72, p < 1e-200), and PC2 and latitude (ρ=−0.55, p < 1e-200). The PCA and birth record data also reflect variability in sampling and population density, with high density in Helsinki and Turku contrasting with low density in the northernmost Lapland region (**Figure S1B**). Mean PC1 and PC2 across birth regions closely mirror geographical patterns, apart from Southern Finland (region 1), which projects closer to central Finland than expected geographically; Southern Finland is the most populous region of Finland, containing the capital city of Helsinki, and consequently draws genetic diversity from across the country (**Figure S1B & Figure S1C**). By comparing parent and offspring birthplaces, we show that within a single generation, offspring across Finland tend to move south, e.g. towards Helsinki (Kolmogorov-Smirnov two-sided test between and child’s and mean parents’ latitude: p=8.7e-3, **Figure S2**).

We also assessed genetic divergence across regions in Finland, and identify relatively high levels of regional divergence compared to other European countries, e.g. the UK, Germany, Sweden, and Estonia ^25,35^, with mean F_ST_ between region pairs = 0.001 (**Figure S1D**); these results are consistent with an additional Finnish bottleneck with respect to nearby countries. Regionally across Finland, we identify geographical clusters with high degrees of similarity. For example, Southern Savonia, Northern Karelia, and Northern Savonia (regions 6, 7, and 8, respectively) exhibit high degrees of genetic similarity (**Figure 1C**). We also identify genetic similarity clusters in the southern central regions of Southern Finland, Tavastia, Southern Karelia, and Central Finland (i.e. regions 1, 4, 5, and 9); western coastal regions of Southwest Finland and Ostrobothnia (2 and 10); and northern regions of Northern Ostrobothnia and Lapland (11 and 12).

**Figure 1.**
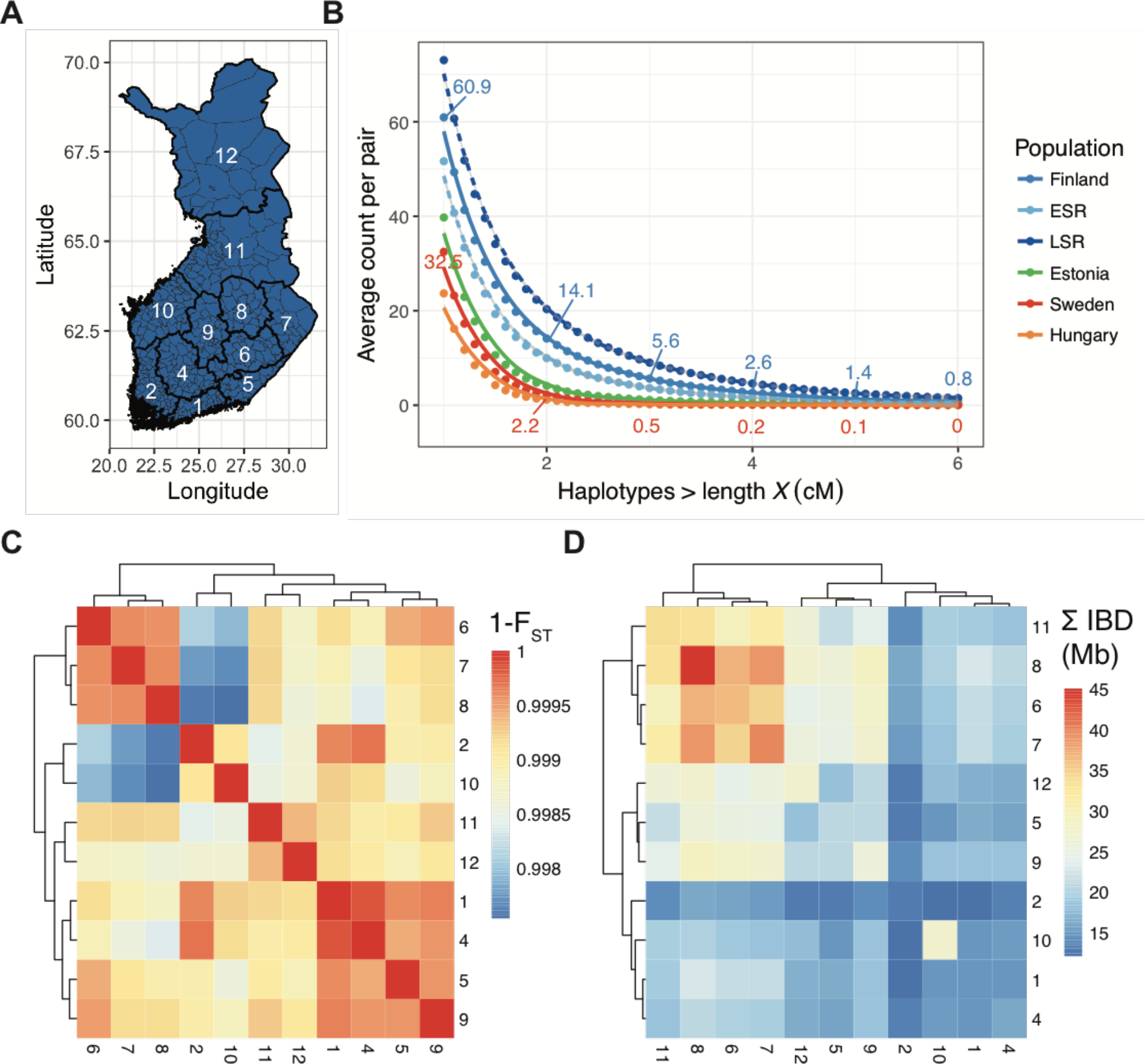
Identity-by-descent (IBD) haplotype sharing and genetic divergence across regions of Finland. A) Regional map of Finland. Region names are shown in **Table S3**. Thin lines within regions represent municipality boundaries. B) Distribution of average pairwise shared IBD segments in Finland (N=7,669), specifically within two birth regions defined previously as having >95% posterior probability of clustering geographically in the ESR (N=428) and LSR (N=592) ^27^, Estonia (N=6,328), Sweden (N=7,863), and Hungary (N=294). All individuals included are unrelated and ancestrally representative of a given region/country. Numbers indicate average pairwise haplotypes shared at 1, 2, 3, 4, and 5 cM in Finland and Sweden. C) Hierarchical clustering of genetic similarity, as measured by 1 - F_ST_ across regions of Finland. D) Hierarchical clustering of cumulative IBD (minimum haplotype ≥ 3 cM) sharing across regions of Finland. C-D) Regions are numbered as in **Table S3**.

### Population bottlenecks in Finland are reflected in identity-by-descent sharing

To better understand the recent population history of Finland, we computed pairwise identity-by-descent (IBD) sharing across all unrelated Finnish pairs of individuals (Methods). We performed hierarchical clustering of cumulative IBD sharing across pairs of individuals within and between regions of Finland, and identified excess sharing in eastern Finland (regions 6, 7, and 8) compared to relatively depleted sharing in southwestern Finland (regions 1, 2, 4, and 10) (**Figure 1D**). Compared to genetic similarity from common variants (**Figure 1C**), haplotype-based clustering is more consistent with historical records that have documented the early vs late settlement areas in southwest and northeast Finland, respectively. Nonetheless, pairwise regional IBD and F_ST_ are highly correlated (Mantel test ρ=0.89, p < 1e-4 with 1,000 Monte Carlo repetitions). Previous work on serial founder effects showed that global genetic divergence increases with geographical distance ^36^, and we recapitulate this finding at the sub-country level within Finland (**Figure S3B**); we also identified decaying IBD sharing with increasing geographical distance within Finland (**Figure S3A**).

Because Finland historically has shared trade, language, and migration with neighboring countries and/or regions, including Sweden, Estonia, and St. Petersburg, Russia, we compared the relative level of allelic and haplotypic sharing within each population. We also compared these genetic data with individuals from Hungary because although it is geographically distal, it shares common linguistic roots; Finnish is a Uralic language that forms an outgroup to most European languages but is related to Estonian and Hungarian. Comparing pairwise IBD sharing within each of these countries, we find that cumulative IBD sharing between pairs of individuals is significantly greater across pairs of individuals on average in Finland than in Sweden, Estonia, Russia, and Hungary, as expected from the Finnish population bottleneck (cumulative total of tracts ≥ 1 cM in length: *µ*_*Sweden*_ = 22.9 cM vs *µ*_*Finland*_ = 107.0 cM, p<1e-50). Consistent with this observation, the average pair of Finns shares more haplotypes that are also longer than in the countries compared here, with for example 13.3 haplotypes ≥ 2 cM shared in Finland vs an order of magnitude fewer (1.3 haplotypes ≥ 2 cM) in Sweden (**Figure 1B**).

### Recent migration inferences from genetic divergence and IBD sharing

We coupled haplotype sharing between pairs of individuals with municipality- and region-level birth record data to determine relative rates of sharing among fine-scale locations in Finland. We subset pairwise IBD to individuals in which both parents were born within 80 km (~50 miles) of each other. For each analysis, we further subset to pairs of individuals in which at least one individual had municipality-level birth records from within 80 km of a given city, then assessed average pairwise IBD with other individuals across municipalities and regions of Finland. By comparing pairwise sharing from different Finnish cities, we find that IBD sharing is very uneven throughout the country, varying by several-fold, and that different geographical regions exhibit considerable substructure with differential IBD sharing patterns (**Figure 2**). This fine-scale structure is likely driven by multiple bottlenecks, recent migration patterns, and variable population density (e.g. genetic diversity is higher and thus IBD sharing is lower in densely populated Helsinki than many rural areas, as Helsinki ancestors have more diverse origins).

**Figure 2.**
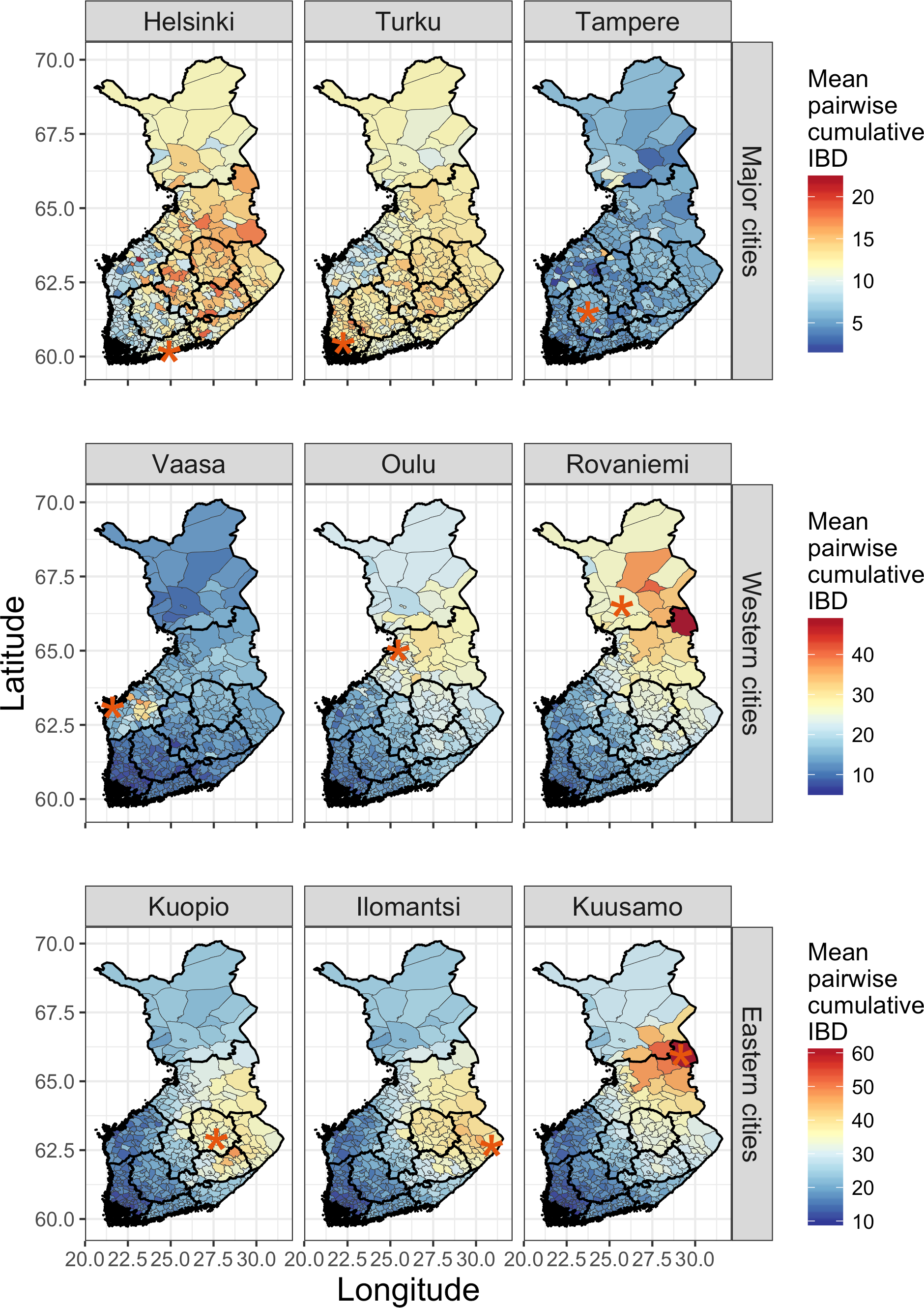
Geographically structured haplotype sharing between pairs of individuals across Finland. We subset to pairs of individuals in which both parents were born within 80 km (~50 miles) of each other. For each panel, we further subset haplotypes from pairs of individuals in which at least one of the individual pairs lives within 80 km of cities indicated by red asterisks. Thinner lines outline municipalities, and thicker lines outline regions. The color shaded in each municipality indicates the weighted mean of cumulative IBD sharing for haplotypes ≥ 3 cM. For each city, the number of unique individuals with both parents from within an 80 km radius and total pairwise comparisons across Finland is as follows: N=152 in Helsinki, 677,844 total pairwise comparisons; N=227 in Turku, 1,003,794 total pairwise comparisons; N=102 in Tampere, 457,419 total pairwise comparisons; N=50 in Vaasa, 225,525 total pairwise comparisons; N=185 in Oulu, 821,955 total pairwise comparisons; N=13 in Rovaniemi, 58,877 total pairwise comparisons; N=566 in Kuopio, 2,406,915 total pairwise comparisons; N=363 in Ilomantsi, 1,580,502 total pairwise comparisons; N=25 in Kuusamo, 113,075 total pairwise comparisons.

Haplotype sharing is on average lowest when at least one individual lives in a major southern Finnish city (**Figure 2**). Specifically, pairwise haplotype sharing across Finland is relatively low across the country with minimal structure when at least one individual lives in Helsinki, Turku, and Tampere. There is subtle structure among individuals born in Helsinki, a relatively young capital (since 1812), indicated by greater haplotype sharing with eastern Finland than western Finland on average (**Figure 2**); in contrast, individuals from the historical capital of Turku, have more elevated haplotype sharing with nearby southwestern Finland (**Figure 2**). IBD sharing is highest among individuals living in northeastern cities in the late settlement area (e.g. Kuopio, Ilomantsi, or Kuusamo), with more structure evident compared to the cosmopolitan cities and greater sharing in the late settlement areas. Of all cities investigated, Kuusamo shows the most elevated IBD sharing, with ~60 Mb on average shared in haplotypes > 3 cM with nearby individuals, compared to ~5-15 Mb near Helsinki, Turku, and Tampere. IBD sharing among western coastal cities (e.g. Vaasa, Oulu, and Rovaniemi) are intermediate, and show varying patterns of regional haplotype sharing. For example, Vaasa, a bilingual city with mostly Finnish and Swedish speakers surrounded by majority Swedish-speaking municipalities, shows restricted patterns of elevated sharing specifically in Ostrobothnia (region 10). Oulu and Rovaniemi in Northern Ostrobothnia and Lapland (regions 11 and 12), in contrast, show broadly elevated patterns of sharing in the late settlement area and depleted sharing in the early settlement area.

We also utilized the granular birth records to investigate geospatial migration rates (*m*). We used a spatially explicit statistical model to estimate effective migration surfaces (EEMS), which measures effective migration rates from genetic differentiation (i.e. resistance distance) across neighboring demes ^37^. By measuring the genetic distance between evenly spaced demes relative to other pairs of demes across Finland and/or neighboring countries, we inferred locations where migration was uncommon, referred to as migration barriers and depicted in dark orange, and where migration excesses occurred, depicted in blue (**Figure 3C-D**). We find variable migration rates across Finland, many of which are consistent with known historical events (**Figure 3C**). For example, we identify barriers to migration generally separating the early and late settlement area (i.e. between Tampere and Kuopio), as well as into the northernmost Lapland region. In contrast, there is increased migration within Finland in/directly surrounding several coastal cities, including Helsinki, Turku, Vaasa, and Oulu.

**Figure 3.**
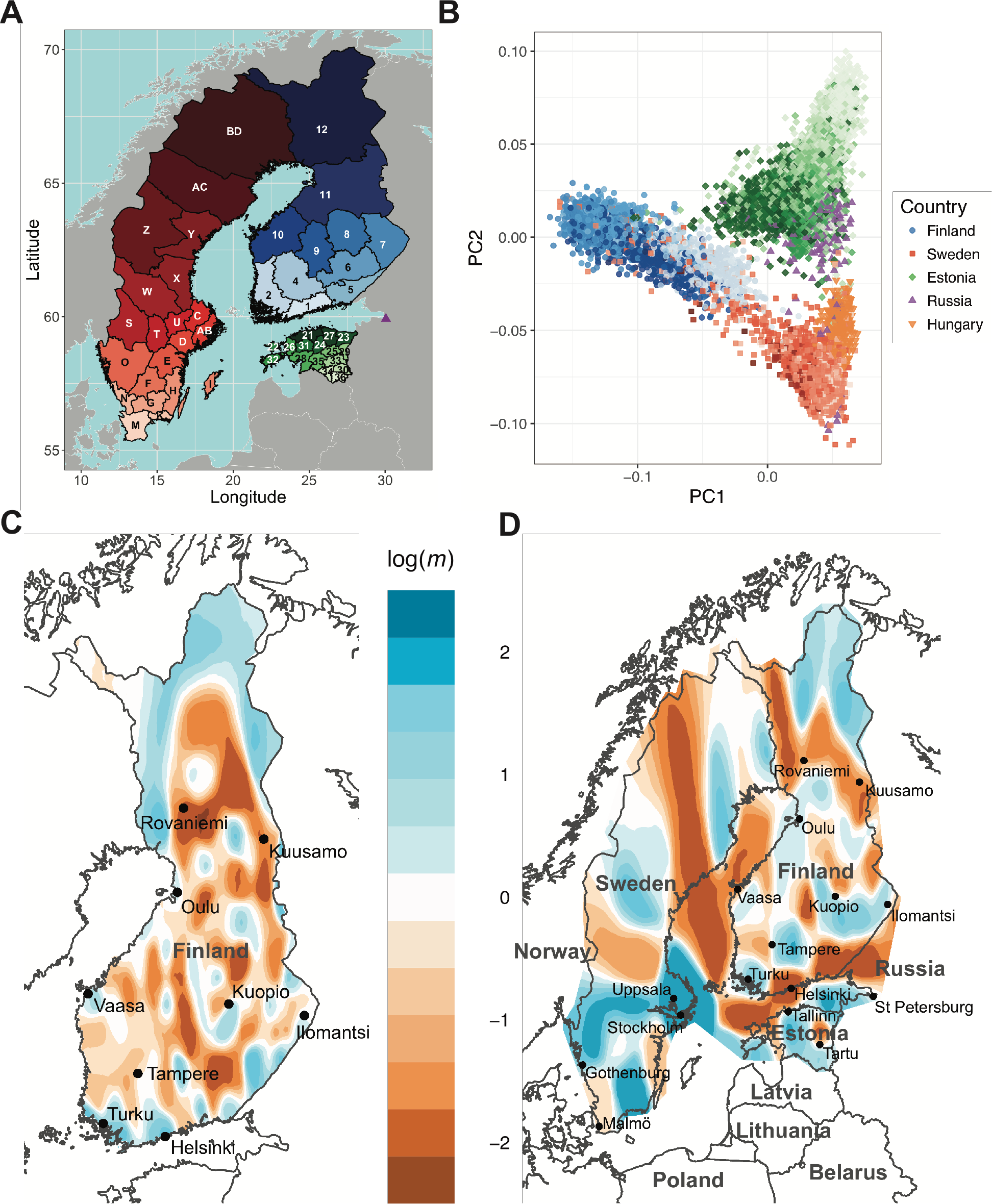
Migration rates and haplotype sharing within Finland and between neighboring countries. A) Map of regional Finnish, Swedish, and Estonian birthplaces. Purple triangle indicates St. Petersburg, Russia. Hungary not shown. Finnish, Swedish, and Estonian region labels are shown in **Table S3**. B) Principal components analysis (PCA) of unrelated individuals, colored by birth region as shown in A) if available or country otherwise. C-D) Migration rates inferred with EEMS. Values and colors indicate inferred rates, for example with +1 (shades of blue) indicating an order of magnitude more migration at a given point on average, and shades of orange indicating migration barriers. C) Migration rates among municipalities in Finland. D) Migration rates within and between Finland, Sweden, Estonia, and St. Petersburg, Russia.

When considering migration rates among individuals with birth records from Finland, Sweden, Estonia, and St. Petersburg, Russia (**Figure 3D**), the major migration routes within Finland remain broadly consistent. For example, a barrier to migration between the early and late settlement regions between Tampere and Kuopio remain, along with a barrier of migration into Lapland. The starkest difference is a barrier to migration along nearly the entire Finnish border (**Figure 3D**), likely due to the absence of some neighboring comparison demes in **Figure 3C** (see also **Figure S5**), indicating little significant migration into Finland in the last 100 generations, consistent with the described patterns of low frequency variation presenting as a bottleneck/isolate. Apart from migration rate inferences along the border, subtle changes within Finland are likely due to additional smoothing because of a larger area over which demes are spread (Methods, **Figure S5**). Migration rates within Sweden are most elevated in southern regions near the largest cities, including Stockholm and Uppsala. As speculated previously ^38^, migration rates are generally elevated within Estonia, but depleted along the west coast and between Tallinn and Tartu; it is also depleted between the Estonia mainland and Finland/Sweden. The strongest barriers to migration in/near Sweden are in the northwest as well as along the northwestern Finnish border separating Finnish Lapland and Sweden, although there are notably few individuals either sampled or living there, resulting in increased noise.

### Fine-scale population differentiation between Finland and nearby countries

We assessed how much sharing occurs within and between regions of Finland and neighboring countries and/or regions, including Sweden, Estonia, St. Petersburg, Russia, and Hungary. PCA recapitulates geographic boundaries and Finnish bottlenecks: PC1 separates Finland from non-Finnish Europeans, and PC2 separates non-Finnish European populations along a cline (**Figure 3B**) ^38,39^. Birth regions also recapitulate expected trends; for example, southern Finns project closer in PCA space with northern Estonians than other regions of either country (**Figure 3B**). Hierarchical clustering of genetic divergence (F_ST_) within and between regions and countries demonstrates that divergence is typically smallest within countries, with the exception Finland and the northernmost Norrbotten region of Sweden that neighbors Finnish Lapland, which cluster together, albeit with the greatest divergence within Finland plus Norrbotten (**Figure S1D**). Taken together with the migration rate analysis, our results suggest that while Norrbotten is most genetically similar to Finnish Lapland, there is still a migration barrier separating these two counties. Individuals from the southwest coastal regions of Finland (regions 1, 2, 10, and 4; i.e. Southern Finland, Southwestern Finland, Ostrobothnia, and Tavastia) are more genetically similar to cosmopolitan Swedes than the rest of the country (**Figure S1D, Figure 3A**). The divergence is greatest (F_ST_ ~ 0.01) between eastern Finland (regions 6, 7, 8; i.e. Southern Savonia, North Karelia, and Northern Savonia) versus Hungary and southern Estonia (regions 30, 34, 36) (**Figure S1D**). The elevated IBD sharing in Finland and the elevated divergence in relation to neighboring countries supports the utility of haplotypes to investigate recent population history as well as IBD mapping to identify rare associations.

### Regional recent effective population size changes over time

Haplotype-sharing also enables a precise assessment of the effective population size of a region. We inferred effective population size changes over recent time across birth regions in Finland using the haplotype-based IBDNe method ^40^. Across all birth regions, we identify a population expansion in the last 50 generations from around 10^3^ to 10^5^ and 10^6^ (**Figure 4, Figure S4**). The region with the largest current effective population size is Southern Finland (region 1, current N_e_=1.3e6), which contains the capital city of Helsinki, closely approximating current census data (current census population ~1.6e6). We inferred that Lapland (region 12), the northernmost and least populated region, had the least growth, with current N_e_=6.9e4 (current census population ~1.8e5). The inferred effective population size is expected to be smaller than the census size because of the census size including multiple generations, variance in reproductive rates, etc ^40^.

When comparing the early versus late settlement areas, we find consistently earlier onset of population expansions in the early settlement area. In the early settlement area, for example, the population began expanding around 30-40 generations ago (circa 760 – 1060 AD, assuming a generation time of 30 years ^41^). In contrast, the late settlement area began expanding between approximately 15-25 generations ago (circa 1210 – 1510 AD), and had lower minimum effective population sizes (**Figure 4**). We also find significant evidence of a geographical cline, wherein populations began expanding earlier in regions further south (ρ=0.79, p=4.2e-3). For example, whereas Southern Finland and Southwestern Finland (regions 1 and 2) began growing ~36 generations ago, the northernmost region of Lapland (region 12) only began growing ~21 generations ago. We also infer larger current effective population sizes in the early rather than late settlement area, consistent with the population density of Finland being higher in the early settlement area. Taken together, the estimation of the regional expansion of the population in conjunction with IBD sharing within and between municipalities provides a clear picture of the history of the population calculated entirely from genetic analysis of the modern Finnish population.

**Figure 4.**
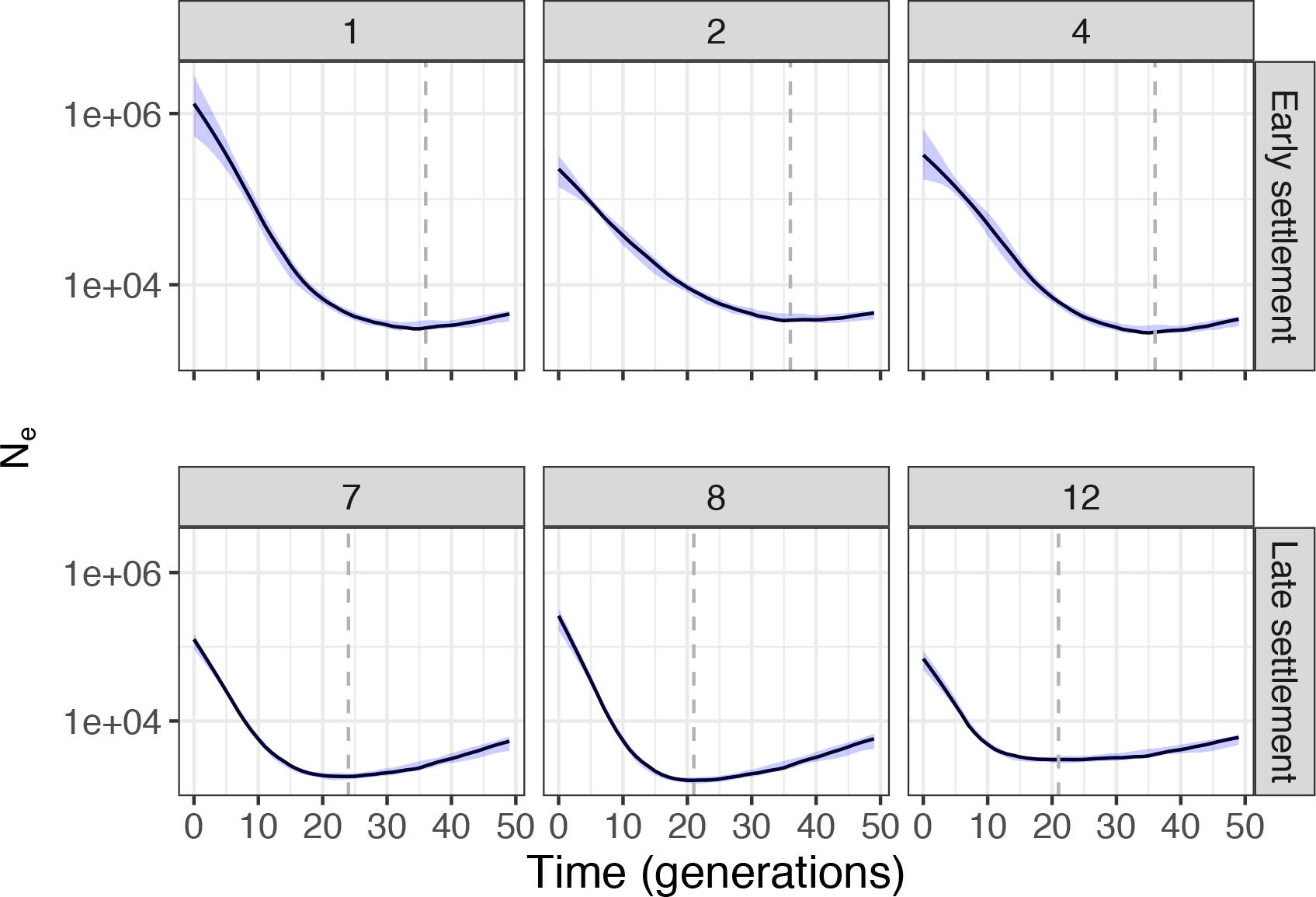
Effective population size over time by birth region in Finland. Representative regions within the early and late settlement areas are numbered as shown in **Table S3**. Dashed lines indicate the time at which the minimum N_e_ over the last 50 generations occurred in each region. Number of individuals in each region are shown in **Figure S4**.

### Haplotype insights into disease

To better understand the utility of IBD sharing for rare variant interpretation, we coupled haplotype tracts with exome sequencing data (Methods). Because previous work in population genetics has suggested that haplotype lengths provide insight into the age of alleles ^42^ and that younger alleles are more likely to be deleterious ^43^, we quantified the extent of haplotype sharing across predicted functional classes of variants and across genotype states. We find as expected that there is generally more haplotype sharing at the rare end of the frequency spectrum (**Figure 5A**). Additionally, we identify greater haplotype sharing in more damaging missense variants than synonymous variants. CpGs disrupt haplotype patterns at the rarest allele frequencies (**Figure 5A, Figure S7**), which is likely a product of mutational recurrence. Haplotype sharing is depleted at the rarest end of the frequency spectrum compared to low frequency variants (~0.1%) in loss-of-function (LoF) and missense constrained genes (**Figure S8**). This depletion may be driven by negative selection against low frequency deleterious variants that are purged prior to reaching more common frequencies ^8,9^ or alternatively because LoF variants are enriched for sequencing error modes ^44^.

**Figure 5.**
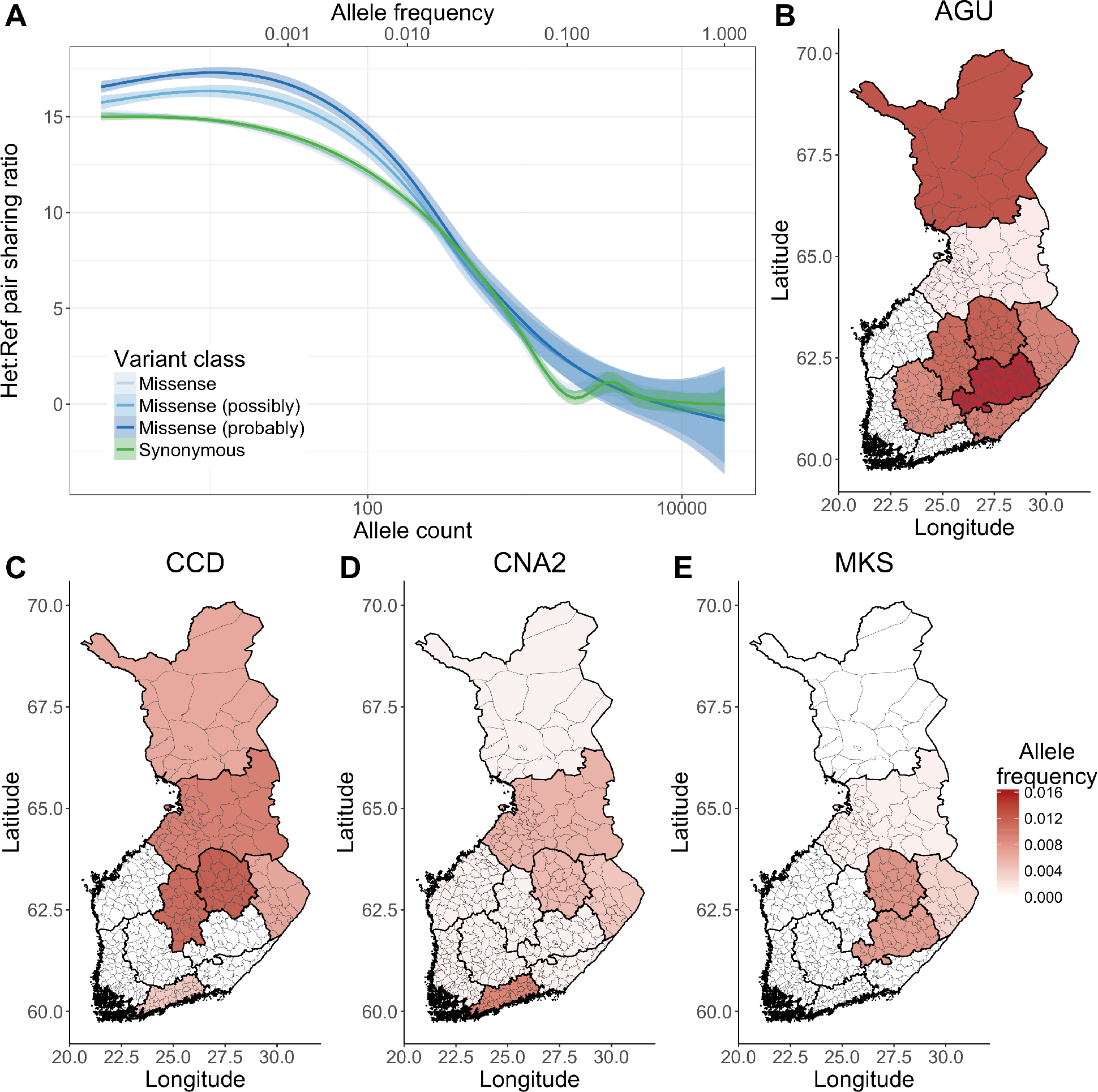
Haplotype sharing enrichment across variant classes and in Finnish heritage diseases. A) Haplotype sharing enrichment among pairs of individuals who are heterozygous versus homozygous reference, excluding CpG variants (Methods). Note that missense (no damaging annotation) and synonymous curves are largely overlapping. B-E) Allele frequency maps for known Finnish heritage disease variants. The same allele frequency scale is included for each of these plots, shown on the bottom right. B) AGU = Aspartylglucosaminuria, C) CNA2 = Cornea plana 2, D) CCD = Congenital chloride diarrhea, and E) MKS = Meckel syndrome. Additional haplotype summaries of these variants are shown in **Table 1**.

We also assessed the overlap of haplotypes for several known disease variants from the Finnish Heritage Disease (FinDis) database (Methods, **Figure 5B-C**). Across the genome, there is a 3% chance that two unselected Finns share a ≥ 1cM haplotype at any position. Considering a set of disease variants with 1% frequency, we first confirmed that indeed homozygous reference individuals (non-carriers) share a haplotype spanning the mutation site at this same background rate. For pairs of individuals who are heterozygous, however, the likelihood of sharing a haplotype ≥ 1 cM is an order of magnitude higher (~30% or higher, **Table 1**). This enrichment of sharing among carriers belies the conceptual framework of IBD mapping, highlighting the power to detect rare, disease-associated loci. We find a significant enrichment of haplotype lengths among pairs of individuals who are heterozygous versus homozygous reference rs386833491 allele (**Figure 5C**). This allele is an in-frame deletion causing congenital chloride diarrhea, and is likely enriched for haplotype sharing beyond the other FinDis variants investigated here because of the regional specificity and origins in the LSR (**Figure 5B**).

**Table 1.** Enrichment of haplotype sharing overlapping FinDis variants. Haplotype enrichment is computed as in **Figure 5** and Methods. the rate of haplotype sharing among pairs of heterozygous individuals per total number of heterozygous pairs relative to homozygous reference pairs. AGU = Aspartylglucosaminuria, CNA2 = Cornea plana 2, CCD = Congenital chloride diarrhea, MKS = Meckel syndrome, FH = familial hypercholesterolemia, T2D = type II diabetes.

| Disease code | gene | rsID | Chr | Pos | Ref | Alt | Freq | Reference pair ratio | Carrier pair ratio | Haplotype enrichment |
| --- | --- | --- | --- | --- | --- | --- | --- | --- | --- | --- |
| AGU | AGA | rs121964904 | 4 | 178359918 | C | G | 0.79% | 0.03 | 0.33 | 10.37 |
| CNA2 | KERA | rs121917858 | 12 | 91449319 | T | C | 0.52% | 0.03 | 0.56 | 16.74 |
| CCD | ACC | rs386833491 | 7 | 107427289 | AACC | A | 0.60% | 0.03 | 0.67 | 21.97 |
| MKS | CC2D2A | rs116358011 | 4 | 15538697 | C | T | 0.30% | 0.03 | 0.28 | 11.31 |

## Discussion

The concept that haplotype tracts assessed from common variant GWAS arrays can provide insight into both population history and rare disease without sequencing data harkens back to the International HapMap Project and before ^11^. While these ideas have been around for decades, their implementation in biobank-scale data is now feasible, and shows promise in isolated populations ^45^. Using data from Finland, we demonstrate that haplotypes provide insight into the evolutionary timeline of greatest interest for this study: recent population history over the past 100 generations and rare, deleterious variants. Coupled with birth record data, haplotype tracts provide deeper insight into fine-scale substructure than common allele approaches alone, including differential sharing within and across coastal and inland municipalities in the early and late settlement areas of Finland.

Finland is particularly amenable for an investigation of recent population history because it has gone through multiple well-documented bottlenecks, has considerable population substructure compared to many other countries ^25–27^, and has a universal health care system with integrated registry information. The relatively high genetic divergence between the early and late settlement areas has been well-documented in prior genetic analyses; we demonstrate much more granular resolution into differential rates of haplotypes across Finland at the level of municipality, for example with several-fold cumulative sharing differences across Finland between major urban southwest cities (e.g. Turku, Helsinki) compared to isolated late settlement areas (e.g. Kuusamo).

The founder effects in Finland have resulted in a massive enrichment of longer haplotypes relative to non-Finnish European neighbors, which depleted genetic diversity overall and increased relatively common deleterious variants with respect to non-Finnish Europeans ^17^. A consequence of these bottleneck signatures is the utility of population-based linkage analysis for discovering deleterious variants at the rare end of the frequency spectrum. Many of the founder mutations contributing to the 36 monogenic diseases in the Finnish Disease Heritage database were originally discovered through family-based linkage analysis ^22^. The emergence of biobank-scale genetic and clinical data enables population-based linkage analysis to discover rare variant associations with previously undiscovered diseases or in populations where risk was previously unrealized, such as a rare orthopedic collagen disorder conferring extreme short stature and dysmorphic features in Puerto Ricans ^45^. Coupling population-based linkage analysis with electronic health records provides a powerful tool for rare disease insights, particularly in populations that have gone through a historical bottleneck. This study demonstrates the utility of haplotype sharing for historical demographic inference and population-based linkage analysis to identify rare variants that confer risk of rare disorders in isolated populations with unified health care registry data, such as Finland.

## Methods

### Genotyping datasets

Finnish samples were genotyped for various projects, all of which have been published previously and most of which were described in ^46^. Briefly, study participants are as follows: European Network for Genetic and Genomic Epidemiology (ENGAGE) Consortium, Myocardial infarction Genetics (MIGen) Exome Array Consortium ^17^, Finrisk (1992, 1997, 2002, and 2007) cohorts, Northern Finland Birth Cohort 1966 (NFBC1966), Corogene controls (which are also from Finrisk), Health 2000 samples from the GenMets study, the Helsinki Birth Cohort Study (HBCS), the Cardiovascular Risk in Young Finns Study (YFS), the Finnish Twin Cohort (FTC). All birth records are from the Finrisk study, which is a superset of several projects. The Finrisk 1997 cohort contains municipality-level birth records (N=3,942), and the 2007 cohort contains region-level birth records (N=5,448), which were genotyped across different projects/arrays (**Table S2**). Swedish samples used here were waves 5 and 6 (Sw5, Sw6) and were genotyped as part of a schizophrenia study ^47^. Swedish genotype data are available upon application from the National Institute of Mental Health (NIMH) Genetics Repository at https://www.nimhgenetics.org/. Estonian samples are from the Estonian Genome Center, University of Tartu (EGCUT) ^38^. Genotyping for individuals from St. Petersburg, Russia was performed as a part of Starvation Study ongoing at the Broad Institute on a cohort previously described in ^48^. Hungarian samples included in the study were genotyped as part of the Hungarian Transdanubian Biobank (HTB) ^49^. Genotyping details and sample sizes are shown in **Table S1**.

### Exome sequencing datasets

Exome sequencing data of Finnish individuals were from multiple studies collected and harmonized as part of Sequencing Initiative Suomi (SISU) study (www.sisuproject.fi, **Table S4**). The Finnish sequence data processing and variant calling has been described previously ^50^. We filtered to exomes with overlapping GWAS data from unrelated individuals this study (N=9,369), as described in “*Haplotypes overlapping exome variants*,” which were primarily from the FINRISK study obtained through dbGaP ^51^. Sample and variant quality control after joint calling differed from that of Rivas et al, 2016 to assess the relationship between rare variation and pairwise haplotype sharing. We filtered to variants present at least twice and excluded variants that failed GATK VQSR quality. See additional information under “*Haplotypes overlapping exome variants*.”

### Phasing and imputation

All Finnish genotypes underwent quality control, phasing, and imputation, as described previously ^46^.

### Principal Components Analysis

We combined best guess genotypes for 43,254 Finnish individuals where variants were imputed with INFO > 0.99 across all arrays, including the Affymetrix Genome-Wide Human SNP 6.0, Illumina Human 370k, 610k, 670k, Core Exome, and OmniExpress arrays. This resulted in ~3.4 million accurately imputed common SNPs across all individuals. From these sites, we performed LD pruning using PLINK v1.90b3f ^52^, keeping SNPs with MAF > 0.05, missingness < 10%, and R^2^ ≤ 0.50 using a window size of 50 SNPs and 5 SNP overlap between windows. PCs were computed across 232,332 sites for all Finnish individuals using flashpca ^53^. We also generated a multi-population dataset of unrelated individuals with birth records where available from Finland, Sweden, Estonia, Hungary, and St. Petersburg, Russia. As before, we extracted best guess Finnish imputed sites with INFO > 0.99. We also filtered to individuals with ≤ 10% missingness, sites with ≤ 10% missingness, MAF ≥ 0.05, and LD R^2^ < 0.5. Because of array heterogeneity, we also filtered to sites on the Illumina Global Screening Array (GSA) to avoid removing all Russian individuals due to high missingness. We then ran PCA with 65,224 sites across N=11,287 individuals.

### Genetic divergence

We computed F_ST_ among geographical regions using PLINK v1.90b3f ^52^. For all analyses, we used the weighted Weir-Cockerham F_ST_ estimate.

### Genetic relatedness

We identified the maximal set of unrelated individuals separated by at least 2 degrees of relatedness using KING v2.0 ^54^ within each population. We identified a maximal unrelated set of: 34,737 Finnish individuals, 7,863 Swedish individuals, 6,328 Estonian individuals, 294 Hungarian individuals, and 210 Russian individuals.

### Haplotype calling

We generated two sets of haplotypes for Finland-only analyses: one for assessing effective population size changes over time using IBDseq ^55^, and another for all other analyses using GERMLINE ^56^. We used IBDSeq rather than GERMLINE for the IBDNe analyses following previous recommendations ^40^ stating that switch errors in estimated haplotypes can cause erroneous haplotype breaks, resulting in spuriously recent time to most recent common ancestor (TMRCA) inferences; IBDseq is less susceptible to these errors since it does not rely on phased data as input. We ran IBDseq on the maximal set of unrelated individuals with birth record data (N=9,008 individuals using 169,306 SNPs). To perform effective population size inferences per region, we subset to haplotypes where both pairs of individuals were born in the same region.

For all other analyses, we first phased all genotype data together using Eagle v2.3.2 ^57^. We then generated haplotype calls using GERMLINE because of its computational tractability at large sample sizes, using the following parameters: -err_hom 0 -err_het 2 - bits 25 -h_extend -haploid. To investigate the decay of IBD tract length, we used a minimum haplotype size of 1 cM (-min_m 1) within each population for unrelated samples with birth record data and/or exome sequencing data. When assessing haplotype sharing across the full set of unrelated genotyped Finns without respect to birth records, we set a minimum haplotype size (-min_m) to 3 cM for computational tractability and reasonable storage sizes. We removed haplotypes that fall partially or fully within centromeres, telomeres, acrocentric short chromosomal arms, heterochromatic regions, clones, and contigs identified in the UCSC hg19 genome “gaps” table.

### Haplotype calling for effective population size analyses

Variants imputed with an info score > 0.99 that intersected across all 6 arrays on which Finnish samples were genotyped (**Table S1**) were included in the haplotype analyses, resulting in 3.4 million accurately imputed common SNPs across 43,254 individuals. High imputation quality best guess genotypes were subsequently filtered to have MAF > 0.05, no indels, and LD R^2^ < 0.5. IBDNe was run across regions of Finland by subsetting to pairs of individuals who were both born in the same region. Demographic analyses included pairwise haplotypes for individuals from the Finrisk 1997 and 2007 cohorts, with the following number of individuals by region: 1,123 in region 1; 1,078 in region 2; 378 in region 4, 224 in region 5; 304 in region 6; 1,581 in region 7; 1,547 in region 8; 225 in region 9; 228 in region 10; 1,697 in region 11; 184 in region 12 (region names as in **Figure 1**).

### Mapping cumulative haplotype sharing

Municipality-level maps of Finland, Sweden, and Estonia were downloaded in R SpatialPolygonsDataFrame (S4) format from http://www.gadm.org/ on 9/14/2015, 4/13/2017, and 7/24/2017, respectively. Pairwise sharing was computed for a maximal unrelated set of individuals (≥ 2^nd^ degree relatives) with municipality- or region-level birth record data (N=8,630 individuals total: N=5,020 with municipality-level data from FR97 and N=3610 with region-level data from FR07). From each city, all pairs where at least one individual had parents born within 80 km of each other and whose mean birth location was within 80 km of the city of interest were included. Municipalities are official and were numbered as described here: https://fi.wikipedia.org/wiki/Luettelo_Suomen_kuntanumeroista, with 3 additional codes: 198 = No home in Finland, 199 = unknown, 200 = abroad. To account for uncertainty when only region-level data was available, even weights were assigned to all municipalities within that region with the sum of the weights equal to 1; in contrast, a single municipality was given a weight of 1 in the municipality-level data.

### Estimating Effective Migration Surfaces (EEMS)

We performed EEMS analysis (Petkova 2015) to estimate migration and diversity relative to geographic distance. We computed genetic dissimilarities for all unrelated pairwise individuals with municipality-level birth record data and both parents born within 80 km, using mean parental latitude and longitude when they differed. We computed pairwise genetic dissimilarities using the *bed2diffs* tool provided with EEMS on the intersected Finnish data with 232,332 SNPs for 2,706 individuals, as well as the intersected Finnish, Swedish, Estonian, and Russian data with 88,080 genotyped SNPs across 10,993 individuals. We set the number of demes to 300 (with fewer actual observed) and adjusted the variances for all proposal distributions of migration, diversity, and degree of freedom parameters such that most were accepted 20-30% of the time and all were accepted 10-40% of the time, per manual recommendations. We increased the number of MCMC iterations, burn-in iterations, and thin iterations until the MCMC converged.

While Finland birth records used in this analysis are at the municipality-level, Swedish and Estonian birth records are at the region-level. Because of differing birth record densities and boundaries in Finland-only versus multi-country analyses, there are differing densities and number of observed demes. When setting nDemes = 300 across Finland, Sweden, Estonia, and St Petersburg, Russia, we observed 110/274 demes. When setting nDemes = 300 across Finland alone, we observed 167/266 demes.

### Haplotypes overlapping exome variants

All analyses of haplotypes paired with exome sequencing data were performed using Hail version 0.1. To map IDs between the genotype and exome sequencing data, we filtered genotype and exome data to variants with at least 1% frequency and less than 10% missingness in each dataset, and subsequently removed individuals with greater than 10% missingness. We intersected these datasets, repeated the same filtering process, and identified 9363 individuals with both data types using the Hail ibd function (minimum pi_hat = 0.95). We assessed haplotype sharing overlapping each SNP used for calling with GERMLINE, and filtered out these overlaps shared at a rate greater than three times the standard deviation above the mean level of sharing (**Figure S6**) to make pairing of exome and haplotype pair data computationally tractable. We overlaid the haplotype data with the exome data using the annotate_variants_table function, and calculated the number of pairs of individuals sharing haplotypes and genotypes for each variant (excluding singletons and variants that failed VQSR filtering) using a custom script in the Hail expression language. Briefly, we determined the set of individuals carrying each genotype and then iterated over the pairs of individuals who share haplotypes, counting cases whether both members of the pair harbored the same genotype. The number of pairs that did not share a given genotype was simply computed as the number of pairs with the genotype 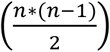 subtracted by the number of pairs that shared the genotype. Variants were subsequently annotated using VEP version 85 using transcripts from Gencode v19 and the LOFTEE plugin (https://github.com/konradjk/loftee; v0.2-28a4843). We then computed the following haplotype enrichment ratio across all exome variants: 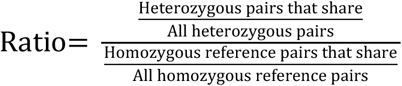.

We stratified haplotype enrichments across allele frequencies and predicted functional variant consequence as well as variants known to cause diseases in the Finnish Disease Heritage database.

## Acknowledgments

Thanks to the participants in the Finnish cohort studies that have made this work possible. Thanks to the sequencing centers at Washington University, the Broad Institute, and the UK10K project for generation and deposition of exome sequencing data from FINRISK and other Finnish cohorts used in the last analyses in this paper. We thank Eimear Kenny and Gillian Belbin for helpful discussions. We also thank Cotton Seed and Tim Poterba for helping scale computational analyses. The Sweden Schizophrenia Study was supported by NIMH R01 MH077139.

## References

1. Lek M, Karczewski KJ, Minikel EV, Samocha KE, Banks E, Fennell T, O’Donnell-Luria AH, Ware JS, Hill AJ, Cummings BB, Tukiainen T, Birnbaum DP, Kosmicki JA, Duncan LE, Estrada K, Zhao F, Zou J, Pierce-Hoffman E, Berghout J, Cooper DN, Deflaux N, DePristo M, Do R, Flannick J, Fromer M, Gauthier L, Goldstein J, Gupta N, Howrigan D, Kiezun A, Kurki MI, Moonshine AL, Natarajan P, Orozco L, Peloso GM, Poplin R, Rivas MA, Ruano-Rubio V, Rose SA, Ruderfer DM, Shakir K, Stenson PD, Stevens C, Thomas BP, Tiao G, Tusie-Luna MT, Weisburd B, Won H, Yu D, Altshuler DM, Ardissino D, Boehnke M, Danesh J, Donnelly S, Elosua R, Florez JC, Gabriel SB, Getz G, Glatt SJ, Hultman CM, Kathiresan S, Laakso M, McCarroll S, McCarthy MI, McGovern D, McPherson R, Neale BM, Palotie A, Purcell SM, Saleheen D, Scharf JM, Sklar P, Sullivan PF, Tuomilehto J, Tsuang MT, Watkins HC, Wilson JG, Daly MJ, MacArthur DG (2016) Analysis of protein-coding genetic variation in 60,706 humans. Nature 536:285–291

2. Rasmussen MD, Hubisz MJ, Gronau I, Siepel A (2014) Genome-Wide Inference of Ancestral Recombination Graphs. PLoS Genetics 10

3. Visscher PM, Wray NR, Zhang Q, Sklar P, McCarthy MI, Brown MA, Yang J (2017) 10 Years of GWAS Discovery: Biology, Function, and Translation. The American Journal of Human Genetics 101:5–22

4. Mathieson I, McVean G (2012) Differential confounding of rare and common variants in spatially structured populations. Nature Genetics 44:243–246

5. Fu W,O’Connor TD, Jun G, Kang HM, Abecasis G, Leal SM, Gabriel S, Altshuler D, Shendure J, Nickerson DA, Bamshad MJ, NHLBI Exome Sequencing Project, Akey JM (2012) Analysis of 6,515 exomes reveals the recent origin of most human protein-coding variants. Nature 493:216–220

6. Kiezun A, Pulit SL, Francioli LC, van Dijk F, Swertz M, Boomsma DI, van Duijn CM, Slagboom PE, van Ommen GJB, Wijmenga C, de Bakker PIW, Sunyaev SR (2013) Deleterious Alleles in the Human Genome Are on Average Younger Than Neutral Alleles of the Same Frequency. PLoS Genetics 9:1–12

7. Zuk O, Schaffner SF, Samocha K, Do R, Hechter E, Kathiresan S, Daly MJ, Neale BM, Sunyaev SR, Lander ES (2014) Searching for missing heritability: designing rare variant association studies. Proceedings of the National Academy of Sciences of the United States of America 111:E455–E464

8. Henn BM, Botigué LR, Peischl S, Dupanloup I, Lipatov M, Maples BK, Martin AR, Musharoff S, Cann H, Snyder MP, Excoffier L, Kidd JM, Bustamante CD (2016) Distance from sub-Saharan Africa predicts mutational load in diverse human genomes. Proceedings of the National Academy of Sciences of the United States of America 113:E440–E449

9. Lohmueller KE (2014) The Impact of Population Demography and Selection on the Genetic Architecture of Complex Traits. PLoS Genetics 10

10. Gutenkunst RN, Hernandez RD, Williamson SH, Bustamante CD (2009) Inferring the joint demographic history of multiple populations from multidimensional SNP frequency data. PLoS genetics 5:e1000695

11. Gibbs RA, Belmont JW, Hardenbol P, Willis TD, Yu FL, Yang HM, Ch’ang L-Y, Huang W, Liu B, Shen Y (2003) The international HapMap project.

12. Bonnen PE, Pe’er I, Plenge RM, Salit J, Lowe JK, Shapero MH, Lifton RP, Breslow JL, Daly MJ, Reich DE, Jones KW, Stoffel M, Altshuler D, Friedman JM (2006) Evaluating potential for whole-genome studies in Kosrae, an isolated population in Micronesia. Nature genetics 38:214–217

13. Sajantila A, Salem AH, Savolainen P, Bauer K, Gierig C, Pääbo S (1996) Paternal and maternal DNA lineages reveal a bottleneck in the founding of the Finnish population. Proceedings of the National Academy of Sciences of the United States of America 93:12035–12039

14. Peltonen L, Peltonen L, Palotie A, Palotie A, Lange K, Lange K (2000) Use of population isolates for mapping complex traits. Nature reviews. Genetics 1:182–190

15. Palo JU, Ulmanen I, Lukka M, Ellonen P, Sajantila A (2009) Genetic markers and population history: Finland revisited. European journal of human genetics: EJHG 17:1336–1346

16. Wang SR, Agarwala V, Flannick J, Chiang CWK, Altshuler D, Hirschhorn JN (2014) Simulation of finnish population history, guided by empirical genetic data, to assess power of rare-variant tests in Finland. American Journal of Human Genetics 94:710–720

17. Lim ET, Würtz P, Havulinna AS, Palta P, Tukiainen T, Rehnström K, Esko T, Mägi R, Inouye M, Lappalainen T, Chan Y, Salem RM, Lek M, Flannick J, Sim X, Manning A, Ladenvall C, Bumpstead S, Hämäläinen E, Aalto K, Maksimow M, Salmi M, Blankenberg S, Ardissino D, Shah S, Horne B, McPherson R, Hovingh GK, Reilly MP, Watkins H, Goel A, Farrall M, Girelli D, Reiner AP, Stitziel NO, Kathiresan S, Gabriel S, Barrett JC, Lehtimäki T, Laakso M, Groop L, Kaprio J, Perola M, McCarthy MI, Boehnke M, Altshuler DM, Lindgren CM, Hirschhorn JN, Metspalu A, Freimer NB, Zeller T, Jalkanen S, Koskinen S, Raitakari O, Durbin R, MacArthur DG, Salomaa V, Ripatti S, Daly MJ, Palotie A (2014) Distribution and Medical Impact of Loss-of-Function Variants in the Finnish Founder Population. PLoS Genetics 10

18. Salmela, E (2012) Genetic structure in Finland and Sweden: aspects of population history and gene mapping. PhD Thesis. University of Helsinki.

19. Poznik GD, Xue Y, Mendez FL, Willems TF, Massaia A, Wilson Sayres MA, Ayub Q, McCarthy SA, Narechania A, Kashin S, Chen Y, Banerjee R, Rodriguez-Flores JL, Cerezo M, Shao H, Gymrek M, Malhotra A, Louzada S, Desalle R, Ritchie GRS, Cerveira E, Fitzgerald TW, Garrison E, Marcketta A, Mittelman D, Romanovitch M, Zhang C, Zheng-Bradley X, Abecasis GR, McCarroll SA, Flicek P, Underhill PA, Coin L, Zerbino DR, Yang F, Lee C, Clarke L, Auton A, Erlich Y, Handsaker RE, Bustamante CD, Tyler-Smith C (2016) Punctuated bursts in human male demography inferred from 1,244 worldwide Y-chromosome sequences. Nature Genetics

20. Kittles RA, Perola M, Peltonen L, Bergen AW, Aragon RA, Virkkunen M, Linnoila M, Goldman D, Long JC (1998) Dual origins of Finns revealed by Y chromosome haplotype variation. The American Journal of Human Genetics 62:1171–1179

21. Tallavaara M, Pesonen P, Oinonen M (2010) Prehistoric population history in eastern Fennoscandia. Journal of Archaeological Science 37:251–260

22. Peltonen L, Jalanko A, Varilo T (1999) Molecular genetics the Finnish disease heritage. Human Molecular Genetics 8:1913–1923

23. Stoll G, Pietiläinen OPH, Linder B, Suvisaari J, Brosi C, Hennah W, Leppä V, Torniainen M, Ripatti S, Ala-Mello S, Plöttner O, Rehnström K, Tuulio-Henriksson A, Varilo T, Tallila J, Kristiansson K, Isohanni M, Kaprio J, Eriksson JG, Raitakari OT, Lehtimäki T, Jarvelin M, Salomaa V, Hurles M, Stefansson H, Peltonen L, Sullivan PF, Paunio T, Lönnqvist J, Daly MJ, Fischer U, Freimer NB, Palotie A (2013) Deletion of TOP3β, a component of FMRP-containing mRNPs, contributes to neurodevelopmental disorders. Nature neuroscience 16:1228–1237

24. Lahtinen AM, Havulinna AS, Jula A, Salomaa V, Kontula K (2015) Prevalence and clinical correlates of familial hypercholesterolemia founder mutations in the general population. Atherosclerosis 238:64–69

25. Salmela E, Lappalainen T, Fransson I, Andersen PM, Dahlman-Wright K, Fiebig A, Sistonen P, Savontaus M, Schreiber S, Kere J, Lahermo P (2008) Genome-Wide Analysis of Single Nucleotide Polymorphisms Uncovers Population Structure in Northern Europe. PLoS ONE 3:e3519

26. Jakkula E, Rehnström K, Varilo T, Pietiläinen OPH, Paunio T, Pedersen NL, deFaire U, Järvelin MR, Saharinen J, Freimer N, Ripatti S, Purcell S, Collins A, Daly MJ, Palotie A, Peltonen L (2008) The Genome-wide Patterns of Variation Expose Significant Substructure in a Founder Population. American Journal of Human Genetics 83:787794

27. Kerminen S, Havulinna AS, Hellenthal G, Martin AR, Sarin AP, Perola M, Palotie A, Salomaa V, Daly MJ, Ripatti S, Pirinen M (2017) Fine-Scale Genetic Structure in Finland. G3 (Bethesda) 7:3459–3468

28. Palamara PF, Lencz T, Darvasi A, Pe’er I (2012) Length distributions of identity by descent reveal fine-scale demographic history. American Journal of Human Genetics 91:809–822

29. Browning SR, Thompson EA (2012) Detecting rare variant associations by identity-by-descent mapping in case-control studies. Genetics 190:1521–1531

30. Ralph P, Coop G (2013) The Geography of Recent Genetic Ancestry across Europe. PLoS biology 11:e1001555

31. Lawson DJ, Hellenthal G, Myers S, Falush D (2012) Inference of population structure using dense haplotype data. PLoS Genetics 8:11–17

32. Szpiech ZA, Xu J, Pemberton TJ, Peng W, Zöllner S, Rosenberg NA, Li JZ (2013) Long runs of homozygosity are enriched for deleterious variation. American journal of human genetics 93:90–102

33. Joshi PK, Esko T, Mattsson H, Eklund N, Gandin I, Nutile T, Jackson AU, Schurmann C, Smith AV, Zhang W, Okada Y, Stančáková A, Faul JD, Zhao W, Bartz TM, Concas MP, Franceschini N, Enroth S, Vitart V, Trompet S, Guo X, Chasman DI, O’Connel JR, Corre T, Nongmaithem SS, Chen Y, Mangino M, Ruggiero D, Traglia M, Farmaki A, Kacprowski T, Bjonnes A, van der Spek A, Wu Y, Giri AK, Yanek LR, Wang L, Hofer E, Rietveld CA, McLeod O, Cornelis MC, Pattaro C, Verweij N, Baumbach C, Abdellaoui A, Warren HR, Vuckovic D, Mei H, Bouchard C, Perry JRB, Cappellani S, Mirza SS, Benton MC, Broeckel U, Medland SE, Lind PA, Malerba G, Drong A, Yengo L, Bielak LF, Zhi D, van der Most PJ, Shriner D, Mägi R, Hemani G, Karaderi T, Wang Z, Liu T, Demuth I, Zhao JH, Meng W, Lataniotis L, van der Laan SW, Bradfield JP, Wood AR, Bonnefond A, Ahluwalia TS, Hall LM, Salvi E, Yazar S, Carstensen L, de Haan HG, Abney M, Afzal U, Allison MA, Amin N, Asselbergs FW, Bakker SJL, Barr RG, Baumeister SE, Benjamin DJ, Bergmann S, Boerwinkle E, Bottinger EP, Campbell A, Chakravarti A, Chan Y, Chanock SJ, Chen C, Chen YI, Collins FS, Connell J, Correa A, Cupples LA, Smith GD, Davies G, Dörr M, Ehret G, Ellis SB, Feenstra B, Feitosa MF, Ford I, Fox CS, Frayling TM, Friedrich N, Geller F, Scotland G, Gillham-Nasenya I, Gottesman O, Graff M, Grodstein F, Gu C, Haley C, Hammond CJ, Harris SE, Harris TB, Hastie ND, Heard-Costa NL, Heikkilä K, Hocking LJ, Homuth G, Hottenga J, Huang J, Huffman JE, Hysi PG, Ikram MA, Ingelsson E, Joensuu A, Johansson Å, Jousilahti P, Jukema JW, Kähönen M, Kamatani Y, Kanoni S, Kerr SM, Khan NM, Koellinger P, Koistinen HA, Kooner MK, Kubo M, Kuusisto J, Lahti J, Launer LJ, Lea RA, Lehne B, Lehtimäki T, Liewald DCM, Lind L, Loh M, Lokki M, London SJ, Loomis SJ, Loukola A, Lu Y, Lumley T, Lundqvist A, Männistö S, Marques-Vidal P, Masciullo C, Matchan A, Mathias RA, Matsuda K, Meigs JB, Meisinger C, Meitinger T, Menni C, Mentch FD, Mihailov E, Milani L, Montasser ME, Montgomery GW, Morrison A, Myers RH, Nadukuru R, Navarro P, Nelis M, Nieminen MS, Nolte IM, O’Connor GT, Ogunniyi A, Padmanabhan S, Palmas WR, Pankow JS, Patarcic I, Pavani F, Peyser PA, Pietilainen K, Poulter N, Prokopenko I, Ralhan S, Redmond P, Rich SS, Rissanen H, Robino A, Rose LM, Rose R, Sala C, Salako B, Salomaa V, Sarin A, Saxena R, Schmidt H, Scott LJ, Scott WR, Sennblad B, Seshadri S, Sever P, Shrestha S, Smith BH, Smith JA, Soranzo N, Sotoodehnia N, Southam L, Stanton AV, Stathopoulou MG, Strauch K, Strawbridge RJ, Suderman MJ, Tandon N, Tang S, Taylor KD, Tayo BO, Töglhofer AM, Tomaszewski M, Tšernikova N, Tuomilehto J, Uitterlinden AG, Vaidya D, van Hylckama Vlieg A, van Setten J, Vasankari T, Vedantam S, Vlachopoulou E, Vozzi D, Vuoksimaa E, Waldenberger M, Ware EB, Wentworth-Shields W, Whitfield JB, Wild S, Willemsen G, Yajnik CS, Yao J, Zaza G, Zhu X, BioBank Japan Project, Salem RM, Melbye M, Bisgaard H, Samani NJ, Cusi D, Mackey DA, Cooper RS, Froguel P, Pasterkamp G, Grant SFA, Hakonarson H, Ferrucci L, Scott RA, Morris AD, Palmer CNA, Dedoussis G, Deloukas P, Bertram L, Lindenberger U, Berndt SI, Lindgren CM, Timpson NJ, Tönjes A, Munroe PB, Sørensen TIA, Rotimi CN, Arnett DK, Oldehinkel AJ, Kardia SLR, Balkau B, Gambaro G, Morris AP, Eriksson JG, Wright MJ, Martin NG, Hunt SC, Starr JM, Deary IJ, Griffiths LR, Tiemeier H, Pirastu N, Kaprio J, Wareham NJ, Pérusse L, Wilson JG, Girotto G, Caulfield MJ, Raitakari O, Boomsma DI, Gieger C, van der Harst P, Hicks AA, Kraft P, Sinisalo J, Knekt P, Johannesson M, Magnusson PKE, Hamsten A, Schmidt R, Borecki IB, Vartiainen E, Becker DM, Bharadwaj D, Mohlke KL, Boehnke M, van Duijn CM, Sanghera DK, Teumer A, Zeggini E, Metspalu A, Gasparini P, Ulivi S, Ober C, Toniolo D, Rudan I, Porteous DJ, Ciullo M, Spector TD, Hayward C, Dupuis J, Loos RJF, Wright AF, Chandak GR, Vollenweider P, Shuldiner AR, Ridker PM, Rotter JI, Sattar N, Gyllensten U, North KE, Pirastu M, Psaty BM, Weir DR, Laakso M, Gudnason V, Takahashi A, Chambers JC, Kooner JS, Strachan DP, Campbell H, Hirschhorn JN, Perola M, Polašek O, Wilson JF (2015) Directional dominance on stature and cognition in diverse human populations. Nature 523:459–462

34. Gamsiz ED, Viscidi EW, Frederick AM, Nagpal S, Sanders SJ, Murtha MT, Schmidt M, Triche EW, Geschwind DH, State MW, Istrail S, Cook EH, Devlin B, Morrow EM (2013) Intellectual disability is associated with increased runs of homozygosity in simplex autism. American Journal of Human Genetics 93:103–109

35. Leslie S, Winney B, Hellenthal G, Davison D, Boumertit A, Day T, Hutnik K, Royrvik EC, Cunliffe B, Lawson DJ, Falush D, Freeman C, Pirinen M, Myers S, Robinson M, Donnelly P, Bodmer W (2015) The fine-scale genetic structure of the British population. Nature 519:309–314

36. Ramachandran S, Deshpande O, Roseman CC, Rosenberg NA, Feldman MW, Cavalli-Sforza LL (2005) Support from the relationship of genetic and geographic distance in human populations for a serial founder effect originating in Africa. Proceedings of the National Academy of Sciences of the United States of America 102:15942–15947

37. Petkova D, Novembre J, Stephens M (2015) Visualizing spatial population structure with estimated effective migration surfaces. Nature Genetics 48:94–100

38. Haller T, Leitsalu L, Fischer K, Nuotio M, Esko T, Boomsma DI, Kyvik KO, Spector TD, Perola M, Metspalu A (2017) MixFit: Methodology for Computing Ancestry-Related Genetic Scores at the Individual Level and Its Application to the Estonian and Finnish Population Studies. PLOS ONE 12:e0170325

39. Nelis M, Esko T, Mägi R, Zimprich F, Zimprich A, Toncheva D, Karachanak S, Piskáčková T, Balaščák I, Peltonen L, Jakkula E, Rehnström K, Lathrop M, Heath S, Galan P, Schreiber S, Meitinger T, Pfeufer A, Wichmann H, Melegh B, Polgár N, Toniolo D, Gasparini P, D’Adamo P, Klovins J, Nikitina-Zake L, Kučinskas V, Kasnauskienė J, Lubinski J, Debniak T, Limborska S, Khrunin A, Estivill X, Rabionet R, Marsal S, Julià A, Antonarakis SE, Deutsch S, Borel C, Attar H, Gagnebin M, Macek M, Krawczak M, Remm M, Metspalu A (2009) Genetic Structure of Europeans: A View from the North–East. PLoS ONE 4:e5472

40. Browning SR, Browning BL (2015) Accurate Non-parametric Estimation of Recent Effective Population Size from Segments of Identity by Descent. The American Journal of Human Genetics 97:404–418

41. Tremblay M, Vézina H (2000) New estimates of intergenerational time intervals for the calculation of age and origins of mutations. Am J Hum Genet 66:651–658

42. Sousa V, Hey J (2013) Understanding the origin of species with genome-scale data: modelling gene flow. Nature reviews. Genetics 14:404–414

43. Rasmussen M, Sikora M, Albrechtsen A, Korneliussen TS, Moreno-Mayar JV, Poznik GD, Zollikofer CPE, Ponce de León MS, Allentoft ME, Moltke I, Jónsson H, Valdiosera C, Malhi RS, Orlando L, Bustamante CD, Stafford TW, Meltzer DJ, Nielsen R, Willerslev E (2015) The ancestry and affiliations of Kennewick Man. Nature

44. MacArthur DG, Tyler-Smith C (2010) Loss-of-function variants in the genomes of healthy humans. Human Molecular Genetics 19:R125–R130

45. Belbin GM, Odgis J, Sorokin EP, Yee M-C, Kohli S, Glicksberg BS, Gignoux CR, Wojcik GL, Van Vleck T, Jeff JM (2017) Genetic Identification Of A Common Collagen Disease In Puerto Ricans Via Identity-By-Descent Mapping In A Health System. bioRxiv:141820

46. Surakka I, Horikoshi M, Mägi R, Sarin A, Mahajan A, Lagou V, Marullo L, Ferreira T, Miraglio B, Timonen S, Kettunen J, Pirinen M, Karjalainen J, Thorleifsson G, Hägg S, Hottenga J, Isaacs A, Ladenvall C, Beekman M, Esko T, Ried JS, Nelson CP, Willenborg C, Gustafsson S, Westra H, Blades M, de Craen AJM, de Geus EJ, Deelen J, Grallert H, Hamsten A, Havulinna AS, Hengstenberg C, Houwing-Duistermaat JJ, Hyppönen E, Karssen LC, Lehtimäki T, Lyssenko V, Magnusson PKE, Mihailov E, Müller-Nurasyid M, Mpindi J, Pedersen NL, Penninx BWJH, Perola M, Pers TH, Peters A, Rung J, Smit JH, Steinthorsdottir V, Tobin MD, Tsernikova N, van Leeuwen EM, Viikari JS, Willems SM, Willemsen G, Schunkert H, Erdmann J, Samani NJ, Kaprio J, Lind L, Gieger C, Metspalu A, Slagboom PE, Groop L, van Duijn CM, Eriksson JG, Jula A, Salomaa V, Boomsma DI, Power C, Raitakari OT, Ingelsson E, Järvelin M, Thorsteinsdottir U, Franke L, Ikonen E, Kallioniemi O, Pietiäinen V, Lindgren CM, Stefansson K, Palotie A, McCarthy MI, Morris AP, Prokopenko I, Ripatti S (2015) The impact of low-frequency and rare variants on lipid levels. Nature Genetics 47:589–597

47. Ripke S, O’Dushlaine C, Chambert K, Moran JL, Kähler AK, Akterin S, Bergen SE, Collins AL, Crowley JJ, Fromer M, Kim Y, Lee SH, Magnusson PKE, Sanchez N, Stahl EA, Williams S, Wray NR, Xia K, Bettella F, Borglum AD, Bulik-Sullivan BK, Cormican P, Craddock N, de Leeuw C, Durmishi N, Gill M, Golimbet V, Hamshere ML, Holmans P, Hougaard DM, Kendler KS, Lin K, Morris DW, Mors O, Mortensen PB, Neale BM, O’Neill FA, Owen MJ, Milovancevic MP, Posthuma D, Powell J, Richards AL, Riley BP, Ruderfer D, Rujescu D, Sigurdsson E, Silagadze T, Smit AB, Stefansson H, Steinberg S, Suvisaari J, Tosato S, Verhage M, Walters JT, Levinson DF, Gejman PV, Kendler KS, Laurent C, Mowry BJ, O’Donovan MC, Owen MJ, Pulver AE, Riley BP, Schwab SG, Wildenauer DB, Dudbridge F, Holmans P, Shi J, Albus M, Alexander M, Campion D, Cohen D, Dikeos D, Duan J, Eichhammer P, Godard S, Hansen M, Lerer FB, Liang K, Maier W, Mallet J, Nertney DA, Nestadt G, Norton N, O’Neill FA, Papadimitriou GN, Ribble R, Sanders AR, Silverman JM, Walsh D, Williams NM, Wormley B, Arranz MJ, Bakker S, Bender S, Bramon E, Collier D, Crespo-Facorro B, Hall J, Iyegbe C, Jablensky A, Kahn RS, Kalaydjieva L, Lawrie S, Lewis CM, Lin K, Linszen DH, Mata I, McIntosh A, Murray RM, Ophoff RA, Powell J, Rujescu D, Van Os J, Walshe M, Weisbrod M, Wiersma D, Donnelly P, Barroso I, Blackwell JM, Bramon E, Brown MA, Casas JP, Corvin AP, Deloukas P, Duncanson A, Jankowski J, Markus HS, Mathew CG, Palmer CNA, Plomin R, Rautanen A, Sawcer SJ, Trembath RC, Viswanathan AC, Wood NW, Spencer CCA, Band G, Bellenguez C, Freeman C, Hellenthal G, Giannoulatou E, Pirinen M, Pearson RD, Strange A, Su Z, Vukcevic D, Donnelly P, Langford C, Hunt SE, Edkins S, Gwilliam R, Blackburn H, Bumpstead SJ, Dronov S, Gillman M, Gray E, Hammond N, Jayakumar A, McCann OT, Liddle J, Potter SC, Ravindrarajah R, Ricketts M, Tashakkori-Ghanbaria A, Waller MJ, Weston P, Widaa S, Whittaker P, Barroso I, Deloukas P, Mathew CG, Blackwell JM, Brown MA, Corvin AP, McCarthy MI, Spencer CCA, Bramon E, Corvin AP, O’Donovan MC, Stefansson K, Scolnick E, Purcell S, McCarroll SA, Sklar P, Hultman CM, Sullivan PF (2013) Genome-wide association analysis identifies 13 new risk loci for schizophrenia. Nature Genetics 45:1150–1159

48. Rotar O, Moguchaia E, Boyarinova M, Kolesova E, Khromova N, Freylikhman O, Smolina N, Solntsev V, Kostareva A, Konradi A, Shlyakhto E (2015) Seventy years after the siege of Leningrad. Journal of Hypertension 33:1772–1779

49. Prasad RB, Lessmark A, Almgren P, Kovacs G, Hansson O, Oskolkov N, Vitai M, Ladenvall C, Kovacs P, Fadista J, Lachmann M, Zhou Y, Sonestedt E, Poon W, Wollheim CB, Orho-Melander M, Stumvoll M, Tuomi T, Pääbo S, Koranyi L, Groop L (2016) Excess maternal transmission of variants in the THADA gene to offspring with type 2 diabetes. Diabetologia 59:1702–1713

50. Rivas MA, Graham D, Sulem P, Stevens C, Desch AN, Goyette P, Gudbjartsson D, Jonsdottir I, Thorsteinsdottir U, Degenhardt F, Mucha S, Kurki MI, Li D, D’Amato M, Annese V, Vermeire S, Weersma RK, Halfvarson J, Paavola-Sakki P, Lappalainen M, Lek M, Cummings B, Tukiainen T, Haritunians T, Halme L, Koskinen LLE, Ananthakrishnan AN, Luo Y, Heap GA, Visschedijk MC, Barrett J, de Lange K, Edwards C, Hart A, Hawkey C, Jostins L, Kennedy N, Lamb C, Lee J, Lees C, Mansfield J, Mathew C, Mowatt C, Newman W, Nimmo E, Parkes M, Pollard M, Prescott N, Randall J, Rice D, Satsangi J, Simmons A, Tremelling M, Uhlig H, Wilson D, Abraham C, Achkar JP, Bitton A, Boucher G, Croitoru K, Fleshner P, Glas J, Kugathasan S, Limbergen JV, Milgrom R, Proctor D, Regueiro M, Schumm PL, Sharma Y, Stempak JM, Targan SR, Wang MH, MacArthur DG, Neale BM, Ahmad T, Anderson CA, Brant SR, Duerr RH, Silverberg MS, Cho JH, Palotie A, Saavalainen P, Kontula K, Färkkilä M, McGovern DPB, Franke A, Stefansson K, Rioux JD, Xavier RJ, Daly MJ, Barrett J, de Lane K, Edwards C, Hart A, Hawkey C, Jostins L, Kennedy N, Lamb C, Lee J, Lees C, Mansfield J, Mathew C, Mowatt C, Newman B, Nimmo E, Parkes M, Pollard M, Prescott N, Randall J, Rice D, Satsangi J, Simmons A, Tremelling M, Uhlig H, Wilson D, Abraham C, Achkar JP, Bitton A, Boucher G, Croitoru K, Fleshner P, Glas J, Kugathasan S, Limbergen JV, Milgrom R, Proctor D, Regueiro M, Schumm PL, Sharma Y, Stempak JM, Targan SR, Wang MH (2016) A protein-truncating R179X variant in RNF186 confers protection against ulcerative colitis. Nature Communications 7:12342

51. Borodulin K, Vartiainen E, Peltonen M, Jousilahti P, Juolevi A, Laatikainen T, Mannisto S, Salomaa V, Sundvall J, Puska P (2014) Forty-year trends in cardiovascular risk factors in Finland. The European Journal of Public Health 25:539–546

52. Chang CC, Chow CC, Tellier LC, Vattikuti S, Purcell SM, Lee JJ (2015) Second-generation PLINK: rising to the challenge of larger and richer datasets. GigaScience 4

53. Abraham G, Inouye M (2014) Fast Principal Component Analysis of Large-Scale Genome-Wide Data. PLoS ONE 9:e93766

54. Manichaikul A, Mychaleckyj JC, Rich SS, Daly K, Sale M, Chen WM (2010) Robust relationship inference in genome-wide association studies. Bioinformatics 26:2867–2873

55. Browning BL, Browning SR (2013) Detecting identity by descent and estimating genotype error rates in sequence data. American Journal of Human Genetics 93:840851

56. Gusev A, Lowe JK, Stoffel M, Daly MJ, Altshuler D, Breslow JL, Friedman JM, Pe’er I (2009) Whole population, genome-wide mapping of hidden relatedness. Genome research 19:318–326

57. Loh P, Danecek P, Palamara PF, Fuchsberger C, A Reshef Y, K Finucane H, Schoenherr S, Forer L, McCarthy S, Abecasis GR, Durbin R, L Price A (2016) Reference-based phasing using the Haplotype Reference Consortium panel. Nature Genetics 48:1443–1448

